# Adipocyte Pten Inhibition Improves Metabolic Health Associated with Expanded Lipid Storage Capacity and Reduced Inflammation

**DOI:** 10.64898/2026.06.20.733549

**Authors:** Yumei Zhou, Yubo Wang, Jessica E. Meerson, Zhiyong Cheng, Shihuan Kuang, Feng Yue

**Author notes:** Correspondence: Feng Yue, Department of Animal Sciences, University of Florida, Gainesville, FL 32611, USA.

## Abstract

Adipose tissue dysfunction drives obesity-associated insulin resistance, but whether expanding adipocyte lipid storage can improve metabolic health remains unclear. Here, we generated adipocyte-specific *Pten* knockout mice (*Pten^AKO^*) using *Adipoq*-Cre to determine how chronic Pten loss affects adipose tissue remodeling and systemic metabolism. *Pten^AKO^* mice exhibit increased adiposity and adipocyte hypertrophy under chow and high-fat diet feeding, yet showing lower blood glucose and insulin levels, enhanced insulin sensitivity, and reduced hepatic lipid accumulation during basal growth and diet-induced obesity without systemic metabolic deterioration. Despite lipid enrichment in brown adipose tissue, Pten-deficient adipocytes maintain UCP1 expression, OXPHOS protein abundance, and mitochondrial ultrastructure. Transcriptomic analysis of inguinal white adipose tissue reveals activation of adipogenesis, lipid metabolism, insulin response, oxidative phosphorylation, lipid storage, vascular and extracellular matrix pathways, together with suppression of immune and inflammatory programs. Mechanistically, Pten deficiency increases *Cav1* expression, caveolae abundance, collagen expression, and extracellular matrix remodeling, suggesting coordinated structural adaptation to support adipocyte expansion. These findings demonstrate that adipocyte Pten deficiency promotes metabolically healthy adipose expansion by enhancing lipid storage capacity, preserving adipocyte function, and reducing inflammation.

## INTRODUCTION

Obesity is a major driver of type 2 diabetes and cardiometabolic disease, but its pathogenic impact extends beyond excess fat mass alone [1]. A central feature of obesity-associated metabolic deterioration is adipose tissue dysfunction, characterized by impaired lipid buffering, chronic low-grade inflammation, altered adipokine secretion, and defective insulin action [2, 3]. When adipose tissue can no longer safely accommodate surplus energy, lipids spill over into liver, skeletal muscle, pancreas, and other non-adipose organs, promoting ectopic lipid deposition, systemic insulin resistance, and widespread metabolic dysfunction [4–6]. Thus, a fundamental question in obesity biology is not simply why adipose tissue enlarges, but why its lipid storage capacity eventually becomes inadequate, thereby converting excess energy storage into a driver of disease [7–9].

Recent studies from human and animal models demonstrate that not all obesity is metabolically equivalent [10–12]. Individuals with similar body mass index can display markedly different metabolic phenotypes, ranging from relatively insulin-sensitive “metabolically healthy” obesity to severe insulin resistance and overt metabolic syndrome [11, 13]. Emerging evidence suggests that this divergence is strongly influenced by adipose tissue quality, depot distribution, and expandability [10, 11]. Metabolically healthier obesity is generally associated with greater subcutaneous fat storage capacity, lower visceral and ectopic lipid accumulation, and more preserved adipose remodeling, whereas metabolically unhealthy obesity is linked to visceral fat expansion, adipocyte stress, inflammation, fibrosis, and lipid overflow [10, 14]. Human studies further support this concept by showing that adipocyte hypertrophy in visceral white adipose tissue (vWAT) is more tightly associated with insulin resistance, while hypertrophic subcutaneous white adipose tissue (sWAT) can be comparatively less inflammatory, underscoring the idea that adipocyte storage capacity and depot-specific remodeling are critical determinants of metabolic health [15–17].

Pten is a key negative regulator of the PI3K-AKT signaling pathway and has emerged as an important determinant of adipose tissue growth, insulin responsiveness, and metabolic homeostasis[18–20]. Early studies using global Pten transgenic mouse model reported that Pten positively regulates brown adipose function, energy expenditure, and longevity [21]. Using aP2-Cre mice to delete *Pten* in adipose tissue, it has been found that loss of Pten enhances insulin sensitivity and confers resistance to streptozotocin-induced diabetes, although it did not markedly alter overall adiposity or circulating fatty acid levels [22]. Subsequent work using an inducible adiponectin-rtTA/TRE-Cre model demonstrated that *Pten* deletion specifically in mature adipocytes is sufficient to enhance adipocyte insulin sensitivity, promote adipose tissue expandability, and improve whole-body metabolic homeostasis [23]. In addition, adult-onset deletion of *Pten* through rAAV-Cre delivery to *Pten^flox^* mice further revealed that adipose Pten regulates adipose tissue homeostasis and fat redistribution through a Pten-Leptin-sympathetic loop [24]. Consistent with these findings in mature adipose tissue, reduction of Pten in adipose progenitor cells by siRNA or CRISPR also enhances progenitor proliferation and adipogenic differentiation [25], suggesting that Pten restrains adipose expansion at multiple levels. Together, these studies identify Pten as a critical brake on adipose tissue remodeling and support the concept that modulation of Pten signaling may influence the capacity of adipose tissue to safely store lipid and thereby shape systemic metabolic health.

In the present study, we used a Cre-mediated constitutive *Pten* knockout (KO) model to define how chronic loss of Pten in adipocytes influences adipose tissue growth and systemic metabolism. Specifically, we investigated how constitutive *Pten* deletion affects adipose expansion, lipid storage, depot remodeling, insulin sensitivity, and whole-body metabolic homeostasis in the setting of obesity. By comparing this constitutive model with prior inducible approaches, our study provides an opportunity to determine how sustained Pten deficiency shapes adipose tissue development and function over time, and to test the concept that increasing adipocyte lipid storage capacity can uncouple obesity from its usual metabolic complications.

## RESULTS

### Adipocyte-specific *Pten* deletion induces increased adiposity in young adult mice

To investigate the role of Pten in adipocytes, we generated adipocyte-specific *Pten* knockout mice by crossing *Adipoq-Cre* mice with *Pten^f/f^* mice (hereafter as *Pten^AKO^*) in which exon 5 encoding the phosphatase domain of Pten was flanked by engineered LoxP sites, leading to premature translational stop and generation of a truncated peptide lacking the key functional phosphatase domain (Figure 1A). In this model, *Pten* should be deleted exclusively in adipocytes as *Adipoq*-Cre specifically marks all mature adipocytes [26]. We confirmed the efficient and specific reduction of Pten protein levels in both BAT and WAT by Western blotting (Figure 1B). Notably, Pten was expressed differentially in different fat depots in WT control mice, with higher level in gonadal WAT (gWAT), moderate level in inguinal and anterior WAT (iWAT and aWAT), and lower level in BAT (Figure 1B).

**Figure 1.**
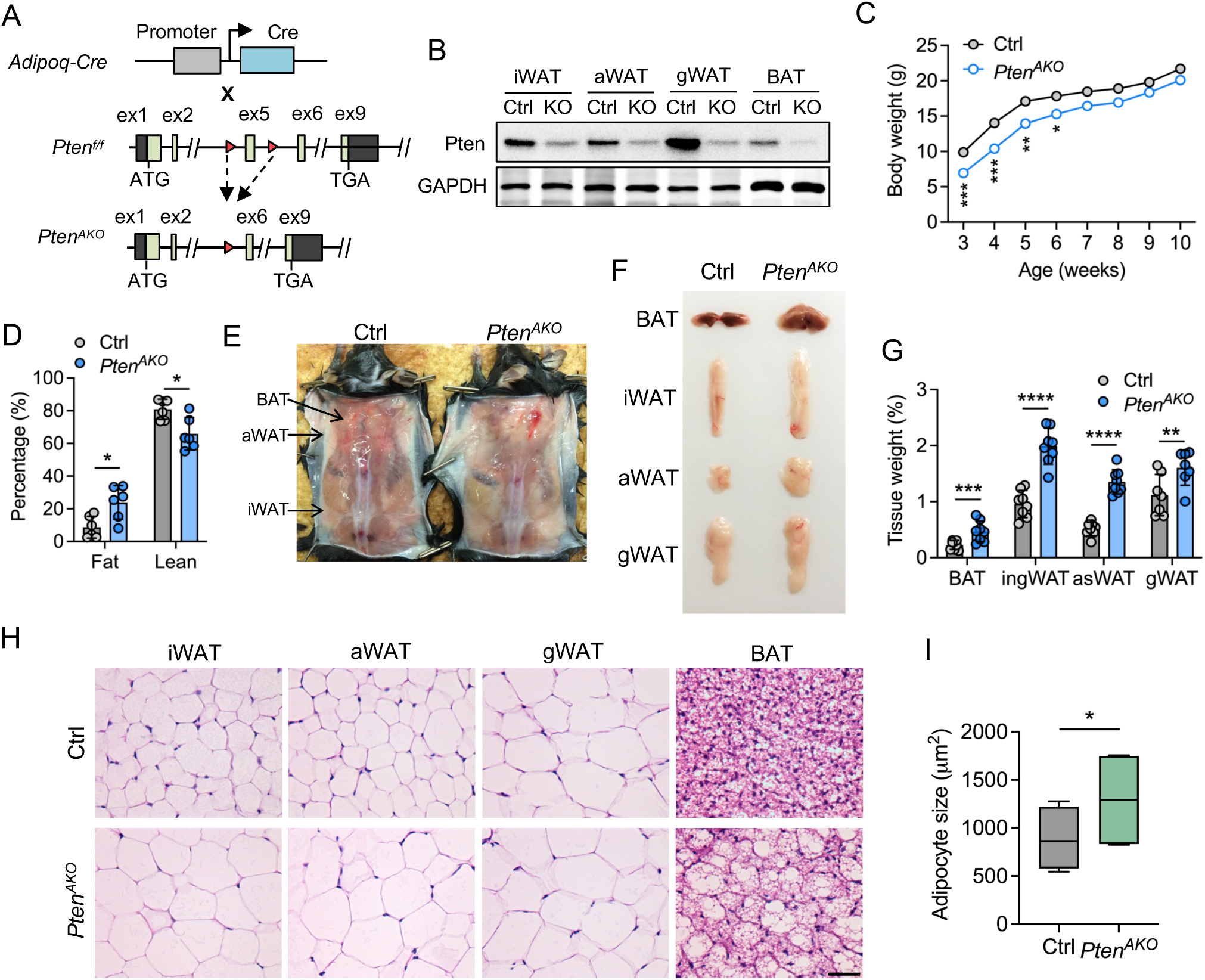
Adipocyte-specific *Pten* deletion induces increased adiposity in young adult mice. *A:* Schematic diagram of the strategy used to generate adipocyte-specific *Pten* knockout (*Pten^AKO^*) mice (ex, exon). *B:* Western blot showing the efficient reduction of Pten protein in WAT and BAT depots of young adult *Pten^AKO^* mice. iWAT, inguinal white adipose tissue; aWAT, anterior subcutaneous white adipose tissue; gWAT, gonadal white adipose tissue; BAT, brown adipose tissue. *C:* Body weight of control (Ctrl) and *Pten^AKO^* mice. *n* = 3-7 mice per group. *D:* Representative image showing the mice and fat from Ctrl and *Pten^AKO^* mice. *E:* Body composition. *n* = 6 mice per group. *F:* Representative image showing the morphology of adipose tissues from Ctrl and *Pten^AKO^* mice. *G:* Quantification of adipose tissue weight normalized to body weight. *n* = 7-8 mice per group. *H:* Representative H&E staining of WAT and BAT depots from control and *Pten^AKO^* mice. Scale bar, 50 µm. *I:* Quantification of adipocyte size of iWAT in Ctrl and *Pten^AKO^* mice. *n* = 4 mice per group. Data are presented as mean ± SEM. \**P* < 0.05, \*\**P* < 0.01, \*\*\**P* < 0.001, and *\*\*\*\*P* < 0.0001 by two-tailed, unpaired Student *t*-test.

The *Pten^AKO^* mice were born at the expected Mendelian ratio and were morphologically indistinguishable from their WT littermates. However, as the *Pten^AKO^* mice grew, they began to have less body weight than control at 3 weeks old and kept at lower body weight until 7 weeks old when comparable body weight was observed with control mice (Figure 1C). Although similar body weight, EchoMRI body composition measurements showed increased total body fat mass and decreased lean mass in *Pten^AKO^* mice compared with control at 2-3 months old under chow diet feeding condition (Figure 1D). Morphologically, the different fat depots of *Pten^AKO^* mice, including BAT, iWAT, asWAT and gWAT, appear to be larger than that of control, with the percentage of tissue weight increased significantly by 106.5%, 107.8%, 164.0%, and 43.5%, respectively (Figures 1E-G). Consistently, the histological analysis revealed the much larger adipocyte size in iWAT, asWAT, and gWAT while more lipid accumulation in BAT (Figures 1 H and I). These data demonstrate that deletion of *Pten* in adipocytes induces increased adiposity with hypertrophic adipocytes in young adult mice.

### Loss of Pten improves systemic metabolism under chow diet feeding

To examine whether the increase of adiposity in the *Pten^AKO^*mice affects their systemic metabolism, we first performed the serum lipid profile. No significant changes were observed in the levels of total triacylglycerol (TG), high-density lipoprotein (HDL) and low-density lipoprotein (LDL) between the *Pten^AKO^* and control mice, whereas the total cholesterol level tends to decrease in the *Pten^AKO^* mice (Figure 2A). In contrast, blood glucose level was significantly reduced in *Pten^AKO^* mice under feeding condition, with 103.0 in *Pten^AKO^* versus 163.7 in control mice (Figure 2B). Remarkably, the serum insulin level in *Pten^AKO^* mice was 3.68-fold lower than the control mice (0.25 versus 0.92) (Figure 2C). We also conducted the glucose tolerance tests (GTT) to determine glucose disposal after intraperitoneal (i.p.) injection of glucose into *Pten^AKO^* and control mice. Lower fasting glucose levels were observed in *Pten^AKO^* mice compared with control mice (Figure 2D). The *Pten^AKO^* mice had lower blood glucose levels than the control littermates after injection of glucose, although the area under curve (AUC) of the *Pten^AKO^*mice were comparable with control mice (Figure 2D). We further performed insulin tolerance tests (ITT) to determine how blood glucose level changes in response to insulin injection (Figure 2E). While the control mice exhibited limited insulin-mediated glucose clearance, a marked decrease of glucose level was observed in *Pten^AKO^*mice (Figure 2E), suggesting the improved insulin sensitivity. Specifically, the area above the curve (AAC) showed a 2.4-fold higher in *Pten^AKO^* mice than control (Figure 2E). Additionally, food intake in *Pten^AKO^* mice was decreased compared to the control (Figure 2F). Together, these results indicate that *Pten* deletion improves insulin sensitivity and systemic metabolism under chow diet feeding.

**Figure 2.**
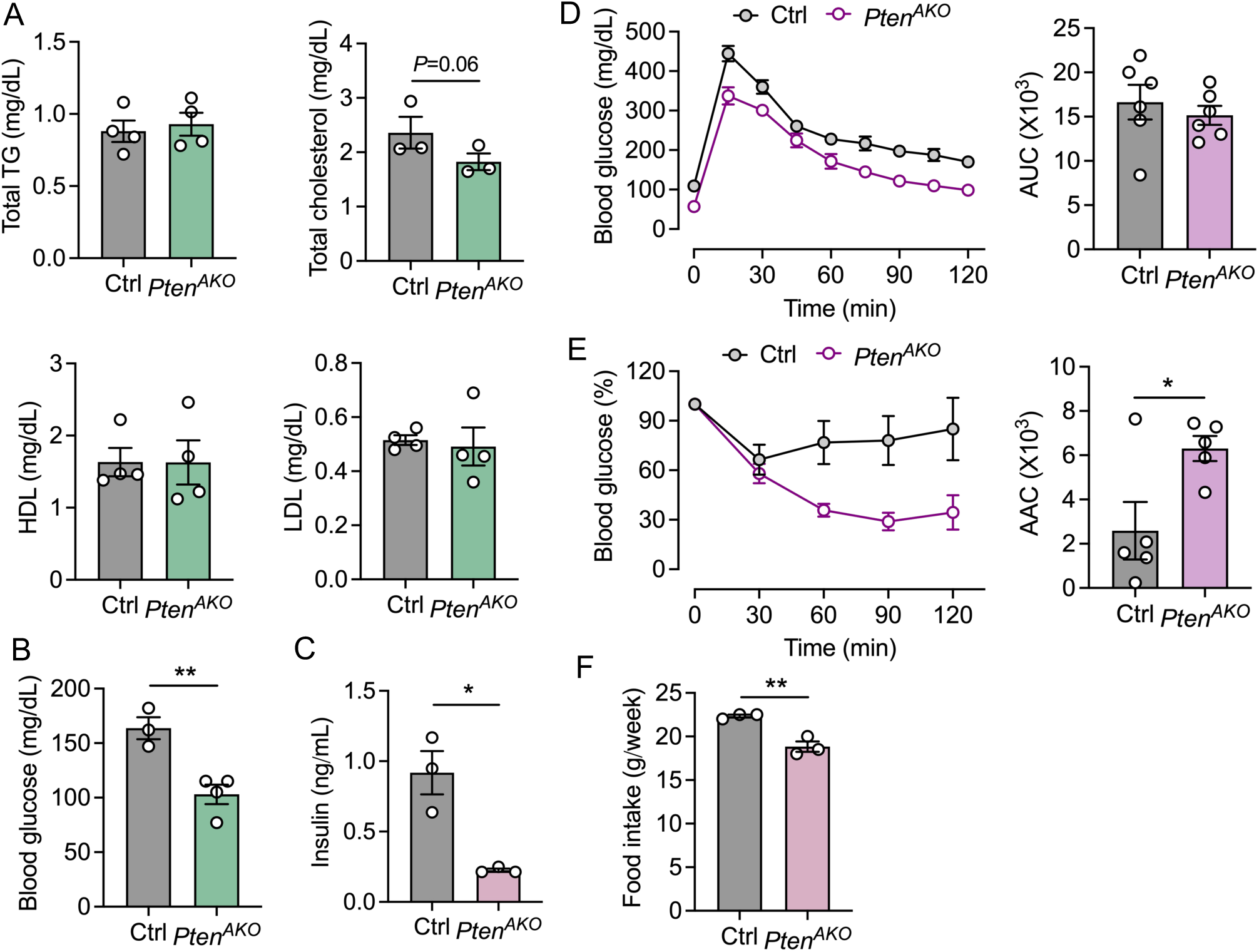
Loss of *Pten* improves systemic metabolism under chow diet feeding. *A:* Serum lipid levels in adult Ctrl and *Pten^AKO^* mice under chow diet feeding condition. *n* = 3-4 mice per group. *B:* Blood glucose level in adult Ctrl and *Pten^AKO^* mice under chow diet feeding condition without fasting. *n* = 3-4 mice per group. *C:* Serum insulin level in adult Ctrl and *Pten^AKO^* mice under chow diet feeding condition. *n* = 3 mice per group. *D:* Glucose tolerance test (GTT) of adult Ctrl and *Pten^AKO^* mice under chow diet feeding condition after 16 hours of fasting. Area under the curve (AUC) was calculated. *n* = 6 mice per group. *E:* Insulin tolerance test (ITT) of adult Ctrl and *Pten^AKO^* mice under chow diet feeding condition after 6 hours of fasting. Area above the curve (AAC) was calculated. *n* = 5 mice per group. *F:* Food intake of adult Ctrl and *Pten^AKO^* mice under chow diet feeding condition. *n* = 3 mice per group. Data are presented as mean ± SEM. \**P* < 0.05 and \*\**P* < 0.01 by two-tailed, unpaired Student *t*-test.

### Pten deficiency leads to excessive expansion of adipose tissue but reduced fat accumulation in liver under HFD feeding

To investigate the role of adipose Pten in mediating animal responses to diet-induced obesity, control and *Pten^AKO^* mice were fed on HFD for 5 months. Surprisingly, there were no discernible differences in body weight between control and *Pten^AKO^* mice at early HFD-feeding (0-10 weeks) (Figure 3A). Although the *Pten^AKO^* mice appear to be fatter, no significant was observed in the body weight after 5 months (Figures 3B and C). However, the fat depots of *Pten^AKO^* mice, including BAT, iWAT, and asWAT appear to be much bigger than that of control (Figures 3D and E), with the percentage of tissue weight increased significantly by 299.7%, 98.2%, and 211.8%, respectively (Figure 3 F). In contrast, the morphology of gWAT was similar between the *Pten^AKO^* and control mice and no significant difference was observed in gWAT mass (Figures 3E and F). In consistent with these observations, the histological analysis revealed the much larger adipocyte size in iWAT, accompanied by crown-like structures and more lipid accumulation in BAT (Figures 3G). Thus, deletion of *Pten* in adipocytes induces excessive expansion of adipose tissue in adult mice under HFD feeding.

**Figure 3.**
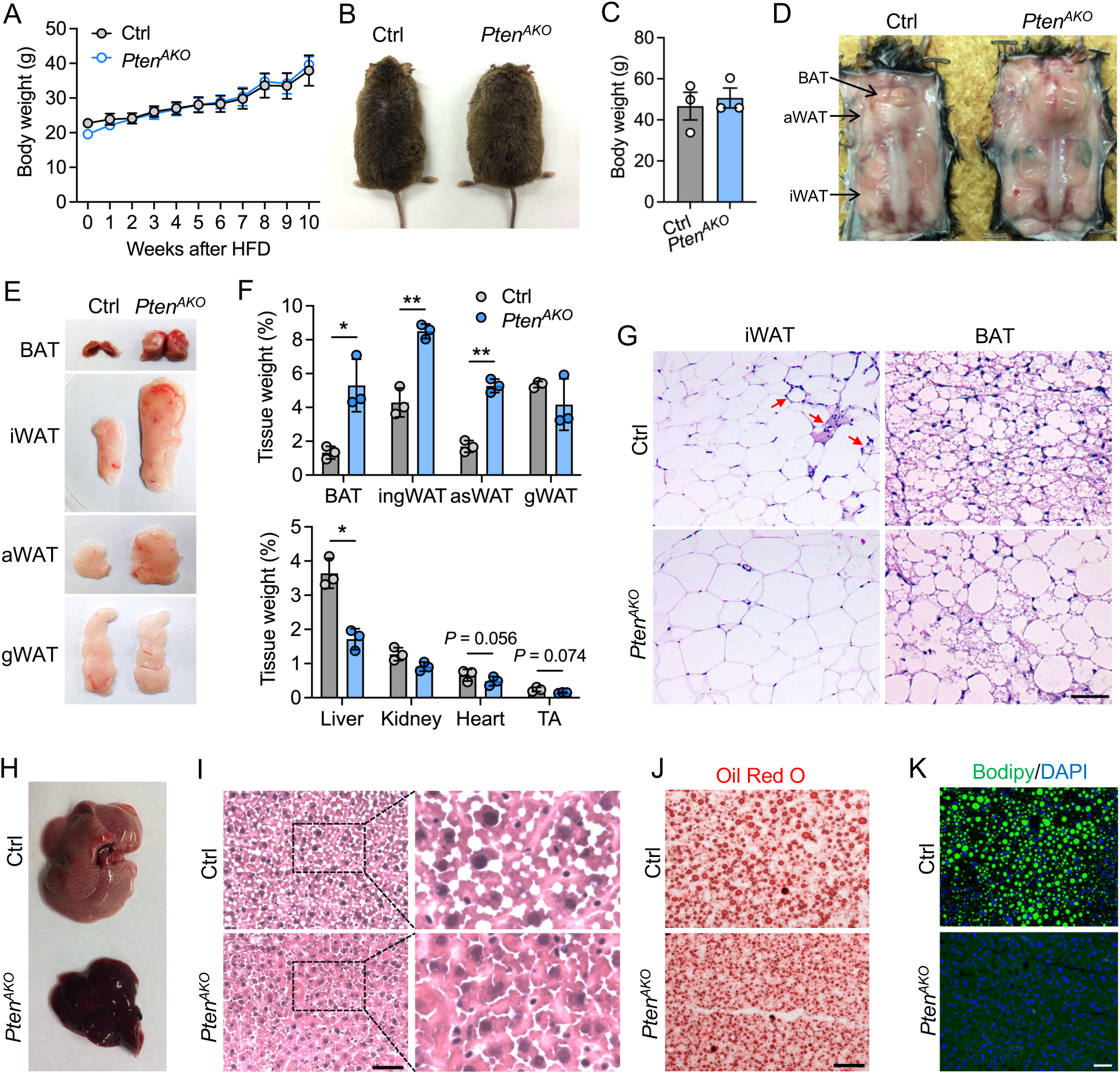
Pten deficiency leads to excessive expansion of adipose tissue but reduced fat accumulation in liver under HFD feeding. *A:* Body weight growth curve of Ctrl and *Pten^AKO^* mice under HFD feeding condition. *n* = 3-4 mice per group. *B:* Representative image of the Ctrl and *Pten^AKO^* mice. *C:* Body weight of Ctrl and *Pten^AKO^* mice under 20 weeks of HFD feeding. *n* = 3 mice per group. *D:* Representative image showing the mice and fat from Ctrl and *Pten^AKO^* mice under 20 weeks of HFD feeding. *E:* Representative image showing the morphology of adipose tissues from Ctrl and *Pten^AKO^* mice under 20 weeks of HFD feeding. *F:* Quantification of tissue weight normalized to body weight under 20 weeks of HFD feeding. *n* = 3 mice per group. *G:* Representative H&E staining of BAT and iWAT from control and *Pten^AKO^* mice under 20 weeks of HFD feeding. Red arrows indicate crown-like structure. *H:* Representative images showing the morphology of liver from Ctrl and *Pten^AKO^* mice under 20 weeks of HFD feeding. *I-K*: Representative H&E (I), Oil Red O (ORO, J), and bodipy staining (K) of liver from control and *Pten^AKO^* mice under 20 weeks of HFD feeding. Scale bar, 50 µm. Data are presented as mean ± SEM. \**P* < 0.05 and \*\**P* < 0.01 by two-tailed, unpaired Student *t*-test.

The drastic increase of subcutaneous fat mass was accompanied with decreased in other tissue mass, including liver, kidney, heart and skeletal muscle (Figure 3F). Interestingly, while the control liver is pale and large, the liver in *Pten^AKO^*was much smaller and red color after HFD feeding (Figure 3H). Further histological analysis revealed more lipid accumulation in control liver, very rare lipid accumulation was observed in *Pten^AKO^* liver (Figure 3I). Consistent results were observed by Oil Red O (ORO) and Bodipy staining (Figures 3J and K). Together, Pten deficiency leads to excessive expansion of adipose tissue but reduced fat accumulation in liver under HFD feeding.

### Improved systemic metabolism and insulin sensitivity of *Pten^AKO^* mice under HFD feeding

We next tested whether the excessive increase of adiposity in the *Pten^AKO^* mice affects their systemic metabolism under HFD feeding. Lipid profile analysis showed that no significant changes were observed in the levels of total TG, LDL and free fatty acids (FFAs) between the *Pten^AKO^* and control mice although the LDL level tends to decrease (Figures 4A and B). Notably, compared to the control, the levels of total cholesterol and HDL were significantly decreased in the *Pten^AKO^* mice, with 3.9 versus 5.7 for cholesterol and 2.1 versus 2.7 for HDL (Figure 4A). Insulin measurements showed that while the control mice developed hyperinsulinemia after HFD feeding, the serum insulin level in *Pten^AKO^* mice was remarkably lower than the control, with 0.166 ng/mL in *Pten^AKO^* mice versus 10.1 ng/mL in control (Figure 4C). In parallel, *Pten^AKO^* mice exhibited significant lower fasting glucose levels compared with control mice, with the average AUC reduced by 63% (Figure 4D). Consistent with the observations in chow diet feeding condition, ITT assay demonstrated a marked decrease of glucose level in *Pten^AKO^* mice after insulin injection, suggesting the improved insulin sensitivity (Figure 4E). Specifically, the quantification of AAC showed a 1.9-fold higher in *Pten^AKO^*mice than control (Figure 4E). These results indicate that the hypertrophic adipocyte induced by Pten deficiency improves adipocyte function under HFD feeding with improved insulin sensitivity and metabolic health.

**Figure 4.**
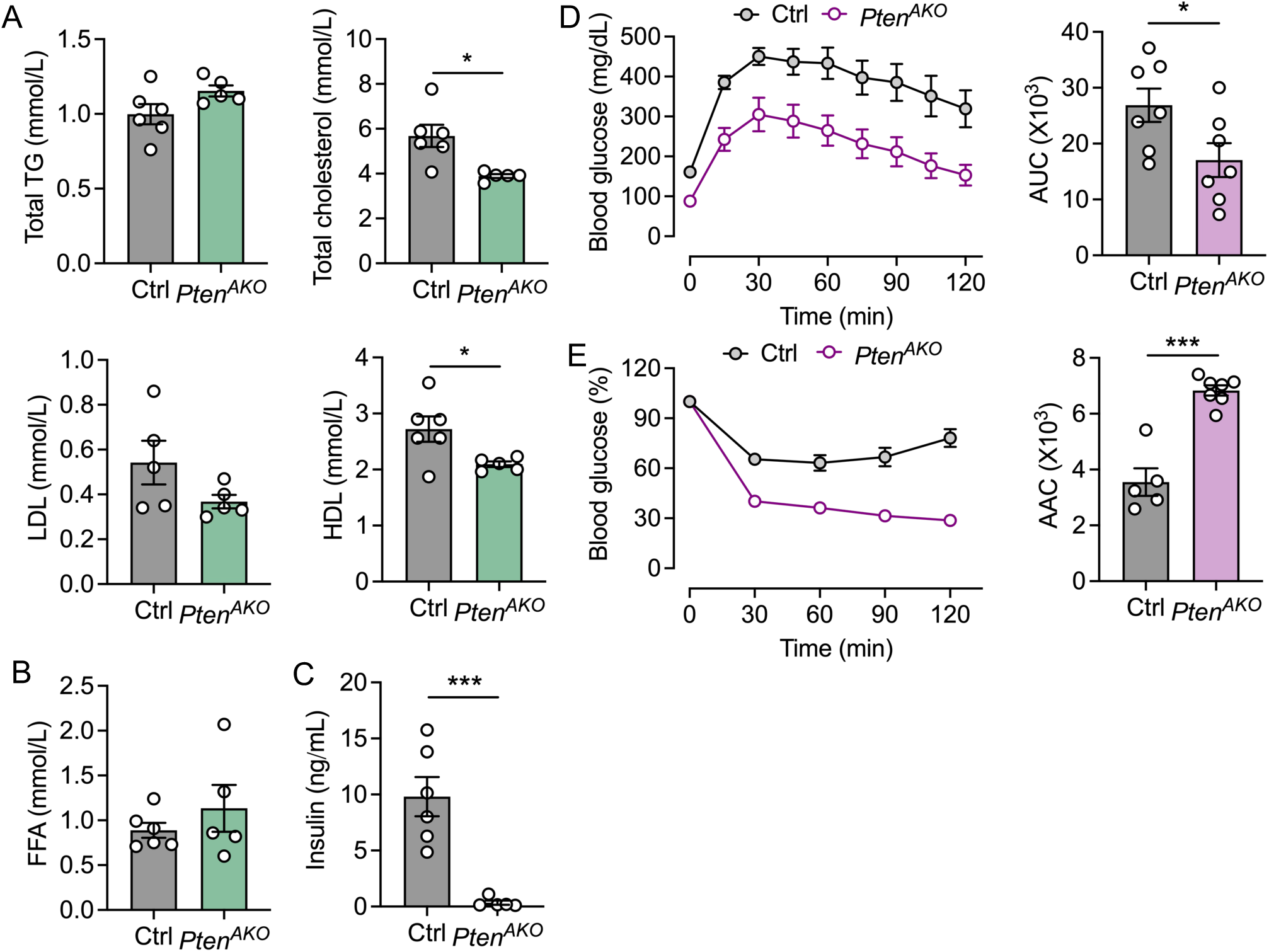
Improved systemic metabolism of *Pten^AKO^* mice under HFD feeding. *A:* Serum lipid levels in adult Ctrl and *Pten^AKO^* mice under HFD feeding condition. Ctrl, *n* = 5-6 mice; *Pten^AKO^*, *n* = 5 mice. *B:* Serum free fatty acid (FFA) level in adult Ctrl and *Pten^AKO^* mice under HFD feeding condition. Ctrl, *n* = 6 mice; *Pten^AKO^*, *n* = 5 mice. *C:* Serum insulin level in adult Ctrl and *Pten^AKO^* mice under 20 weeks of HFD feeding. Ctrl, *n* = 6 mice; *Pten^AKO^*, *n* = 5 mice. *D:* Glucose tolerance test (GTT) of adult Ctrl and *Pten^AKO^* mice under HFD feeding condition after 16 hours of fasting. *n* = 7 mice per group. *E:* Insulin tolerance test (ITT) of adult Ctrl and *Pten^AKO^* mice under HFD feeding condition after 6 hours of fasting. Ctrl, *n* = 5 mice; *Pten^AKO^*, *n* = 7 mice. Data are presented as mean ± SEM. \**P* < 0.05 and \*\*\**P* < 0.001 by two-tailed, unpaired Student *t*-test.

### Hypertrophic *Pten*-deficient adipocytes maintain mitochondrial integrity

Given the excessive lipid storage in *Pten*-deficient adipocytes, we next sought to understand whether the hypertrophy of adipocytes disrupts its mitochondrial function, particularly in BAT. Western blot analysis demonstrated the elimination of Pten protein in *Pten^AKO^* BAT with the markedly activation of phosphorylated AKT (pAKT) (Figures 5A and B). Surprisingly, the expression level of PGC1α was increased in *Pten^AKO^* BAT, while the expression levels of PPARγ, the brown fat marker UCP1 and the adipogenic FABP4 proteins were comparable between *Pten^AKO^* and control mice (Figures 5A and C). Moreover, the expression levels of mitochondrial Oxidative phosphorylation (OXPHOS) complexes were also comparable between *Pten^AKO^*and control mice (Figures 5A and D), suggesting the maintained mitochondrial contents in *Pten*-deficient brown adipocytes. Similar observations were observed in iWAT, in which exhibited activation of pAKT but no changes in OXPHOS complexes (Figures 5E and F). To confirm the expression of UCP1 in *Pten^AKO^* BAT, we further performed immunohistochemical analysis. Apart from the accumulation of large lipid droplets, the *Pten^AKO^* BAT exhibited high intensity of UCP1 staining signal, comparable with the control (Figure 5H). To examine whether the mitochondrial structure was influenced in *Pten^AKO^* BAT, we subjected the BAT to Transmission Electron Microscopy (TEM) analysis. Mitochondria in both control and *Pten^AKO^*BAT exhibited similar size of mitochondria and clear cristae structure (Figure 5I), suggesting the intact integrity of mitochondria. Together, *Pten*-deficient adipocytes maintain mitochondrial content and integrity.

**Figure 5.**
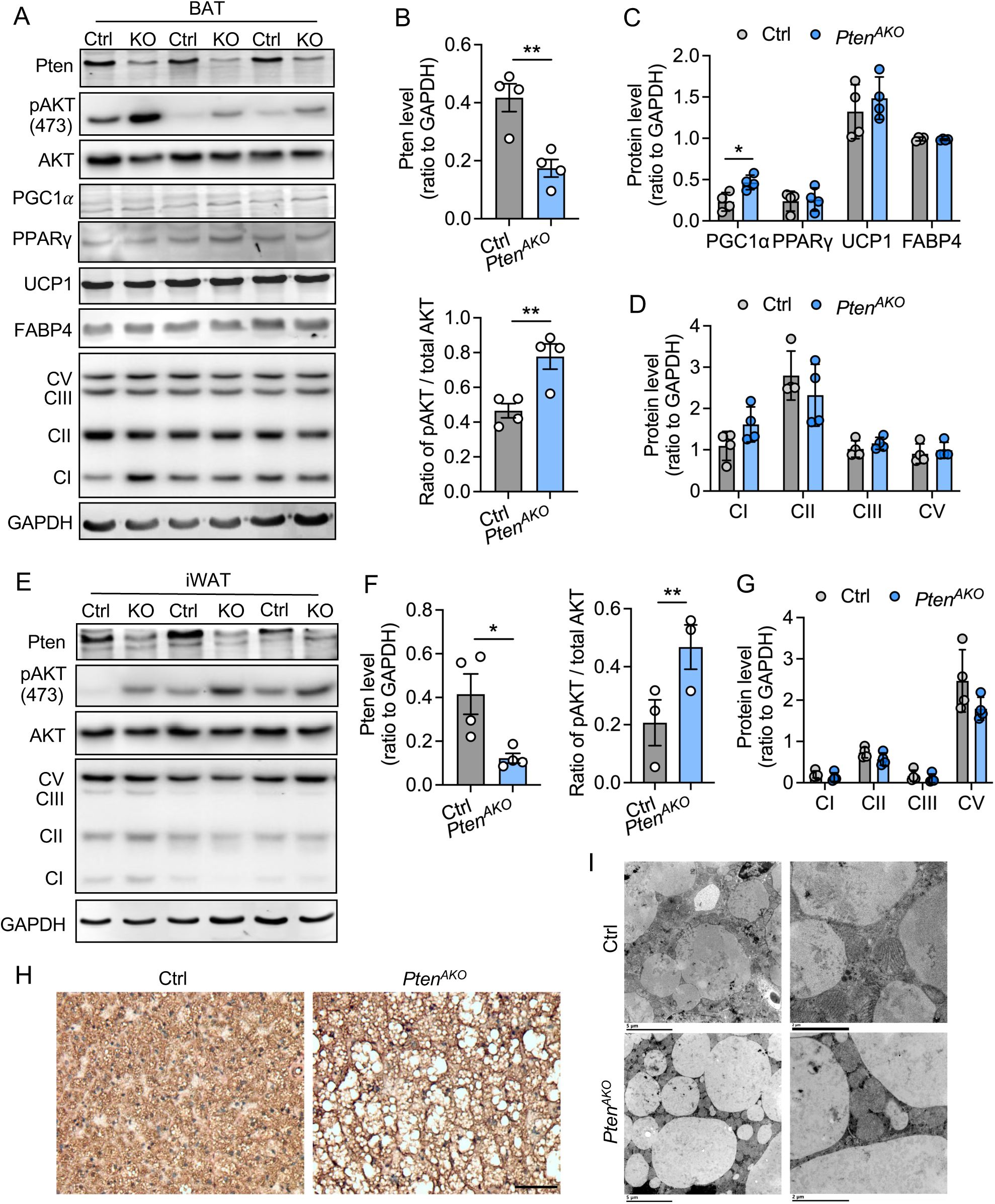
Hypertrophic *Pten*-deficient adipocytes maintain mitochondrial integrity. *A*: Western blot analysis of pAKT and proteins related to lipid and mitochondrial metabolism in BAT of Ctrl and *Pten^AKO^* mice under chow diet feeding condition. *B*: Quantification of Pten and pAKT protein levels shown in *A*. *n* = 4 mice per group. *C*: Quantification of PGC1α, PPARγ, UCP1 and FABP4 protein levels shown in *A*. *n* = 4 mice per group. *D*: Quantification of mitochondrial OXPHOS protein levels shown in *A*. *n* = 4 mice per group. *E*: Western blot analysis of pAKT and mitochondrial OXPHOS proteins in iWAT of Ctrl and *Pten^AKO^* mice under chow diet feeding condition. *F*: Quantification of Pten and pAKT protein levels shown in *E*. *n* = 4 mice per group. *G*: Quantification of mitochondrial OXPHOS protein levels shown in *E*. *n* = 4 mice per group. *H*: Immunohistochemical (IHC) staining of UCP1 in BAT of Ctrl and *Pten^AKO^* mice under chow diet feeding condition. Scale bar, 50 µm. *I*: Transmission electron microscopy (TEM) imaging showing mitochondrial structure in BAT of of Ctrl and *Pten^AKO^* mice. Scale bar, left panel (5 µm), right panel (2 µm). Data are presented as mean ± SEM. \**P* < 0.05 and \*\**P* < 0.01 by two-tailed, unpaired Student *t*-test.

### Microarray analysis reveals the molecular pathways underlying improved adipose tissue function in *Pten^AKO^* mice

To define the molecular pathways underlying the improved adipose tissue function in *Pten^AKO^* mice, we performed microarray analysis using iWAT from adult control and *Pten^AKO^* mice. This analysis identified 1,308 differentially expressed genes (DEGs) in *Pten^AKO^*iWAT compared with controls, including 783 upregulated and 525 downregulated genes (Figure 6A, Supplementary Datasheet 1). Gene Ontology (GO) analysis showed that downregulated genes were strongly enriched for immune-related biological processes, including lymphocyte differentiation, T cell activation, mononuclear cell proliferation, immune effector function, chemotaxis, TNF production, and extrinsic apoptotic signaling (Figure 6B, Supplementary Datasheet 2). In contrast, upregulated genes were enriched for processes associated with cellular respiration, fatty acid metabolism, adipogenesis, triglyceride metabolism, lipid storage, insulin response, extracellular matrix organization, vascular development, and collagen organization (Figure 6C). Notably, genes involved in adipogenesis and insulin responsiveness were increased in *Pten^AKO^* iWAT, including *Slc2a4* (*Glut4*), *Lpl*, *Scd1*, *Adipoq*, *Fabp4*, *Leptin*, *Igf1*, *Insig1*, *Insig2*, and *Xbp1* (Figure 6D).

**Figure 6.**
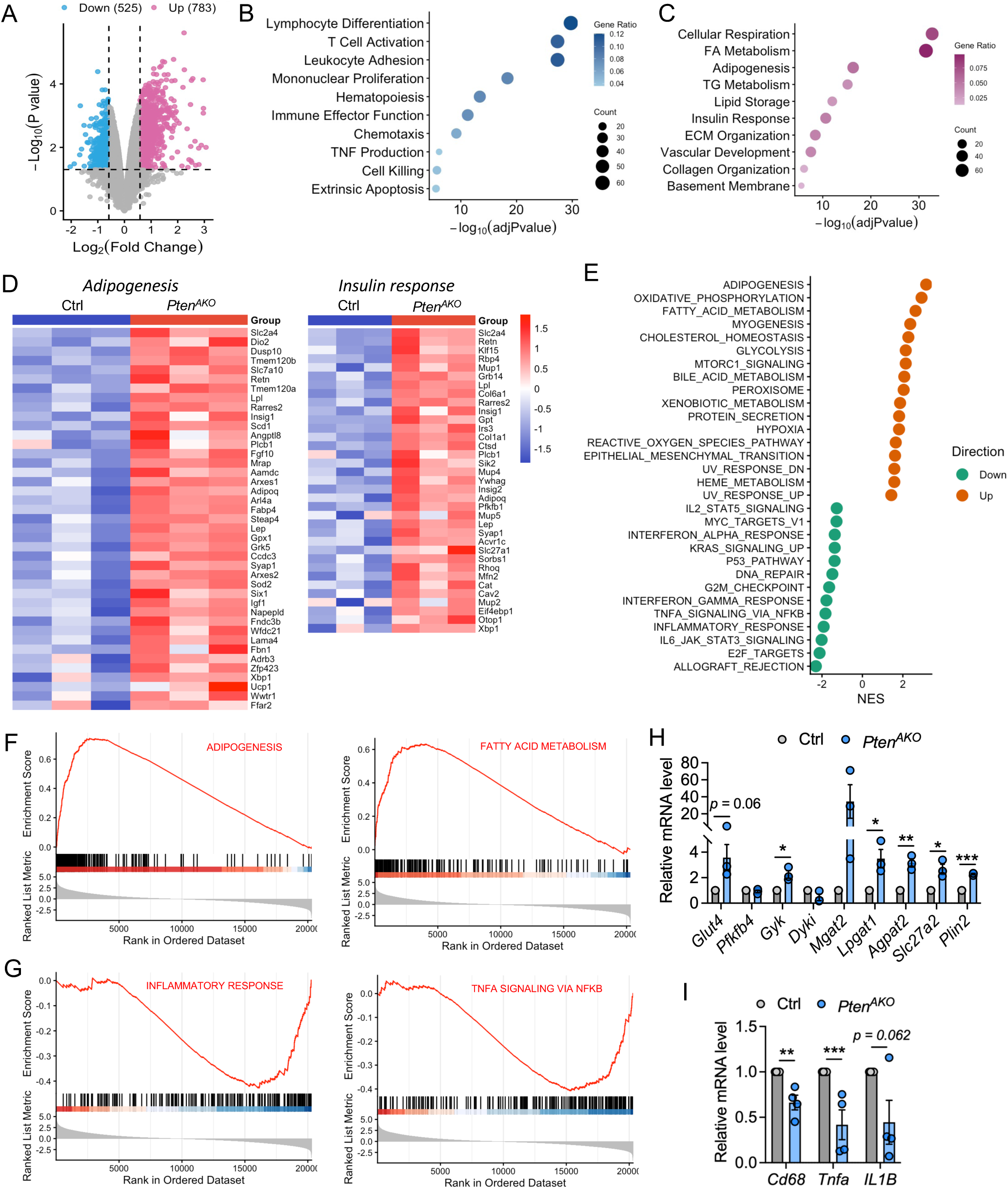
Microarray analysis indicates molecular pathways underlying improved adipose tissue function in *Pten* KO mice. *A*: Volcano plot showing differentially expressed genes (DEGs) between Ctrl and *Pten^AKO^* iWAT. *n* = 3 mice per group. *B and C*: Gene Ontology (GO) enrichment analysis of significantly downregulated (*B*) and upregulated (*C*) genes. *D*: Heatmaps of upregulated genes from microarray analysis related to adipogenesis and insulin response. *E:* Gene Set Enrichment Analysis (GSEA) of significantly upregulated and downregulated genes. *F*: GSEA identified the enrichments of adipogenesis and fatty acid metabolism pathways upregulated in *Pten^AKO^* WAT. *G*: GSEA identified the enrichments of inflammation response and TNFa signaling downregulated in *Pten^AKO^* WAT. *H:* qPCR analysis showing the relative expression of lipid metabolism genes in Ctrl and *Pten^AKO^* iWAT. *n* = 3 mice per group. I: qPCR analysis showing the relative expression of inflammation response genes in Ctrl and *Pten^AKO^* WAT. *n* = 4 mice per group. Data are presented as mean ± SEM. \**P* < 0.05, \*\**P* < 0.01 and \*\*\**P* < 0.001 by two-tailed, unpaired Student *t*-test.

Consistent with the GO analysis, GSEA pathway enrichment analysis of upregulated genes revealed significantly enrichment of adipogenesis, oxidative phosphorylation, fatty acid metabolism, cholesterol homeostasis, glycolysis, peroxisome, xenobiotic metabolism, mTORC1 signaling, and reactive oxygen species pathways (Figure 6E, Supplementary Datasheet 3). In contrast, downregulated genes were enriched for inflammatory response, TNFα signaling, DNA repair, p53 pathway, interferon-α and interferon-γ responses, and IL2–STAT5 signaling pathways (Figure 6E, Supplementary Datasheet 3). GSEA further supported activation of adipogenesis and fatty acid metabolism, together with suppression of TNFα signaling and inflammatory responses in *Pten^AKO^*iWAT (Figures 6F and 6G).

To validate these transcriptomic changes, we performed qPCR analysis of selected genes involved in lipid metabolism and inflammation. The expression of *Glut4*, *Lpgat1*, *Agpat2*, *Slc27a2* and *Plin2* was increased in *Pten^AKO^* iWAT compared with controls (Figure 6H). Conversely, the expression of inflammatory markers, including *Cd68*, *Tnfa*, and *Il1b*, was reduced in *Pten^AKO^* iWAT (Figure 6I). Together, these findings indicate that adipocyte Pten deficiency promotes a transcriptional program associated with enhanced lipid metabolism, adipogenesis, and insulin responsiveness while suppressing inflammatory signaling, providing a molecular basis for improved adipose tissue function in *Pten^AKO^*mice.

### Remodeling of caveolae and ECM facilitates healthy expansion of adipose tissue

We next sought to determine how Pten-deficient adipocytes accommodate expanded lipid storage capacity without apparent adipose dysfunction. Transcriptomic analysis showed increased expression of genes involved in lipid storage, including *Dgat1*, *Dgat2*, *Clstn3*, *Cidec*, *Cd36*, and *Abhd5* (Figure 7A) [27]. Notably, *Cav1*, which encodes caveolin-1, was increased 2.6-fold in *Pten^AKO^* iWAT compared with control iWAT (Figure 7A, Supplemental Datasheet 1). CAV1 is the principal structural component of caveolae, flask-shaped invaginations of the adipocyte plasma membrane that help regulate lipid handling and enable adipocytes to expand safely during lipid accumulation [28–30]. qPCR analysis confirmed the increased expression of *Cav1* in *Pten^AKO^*iWAT (Figure 7B). To determine whether increased *Cav1* expression was associated with structural remodeling of caveolae, we performed transmission electron microscopy analysis of control and *Pten^AKO^* iWAT. Strikingly, *Pten^AKO^*adipocytes displayed a marked increase in the number of caveolae along the plasma membrane compared with control adipocytes (Figure 7C). Quantification showed that caveolae density was increased approximately 2.5-fold in *Pten^AKO^* iWAT, with 15.5 caveolae per membrane area in *Pten^AKO^* adipocytes compared with 6.2 in control adipocytes (Figure 7D). These results suggest that Pten deficiency promotes plasma membrane remodeling that may enhance adipocyte lipid storage capacity.

**Figure 7.**
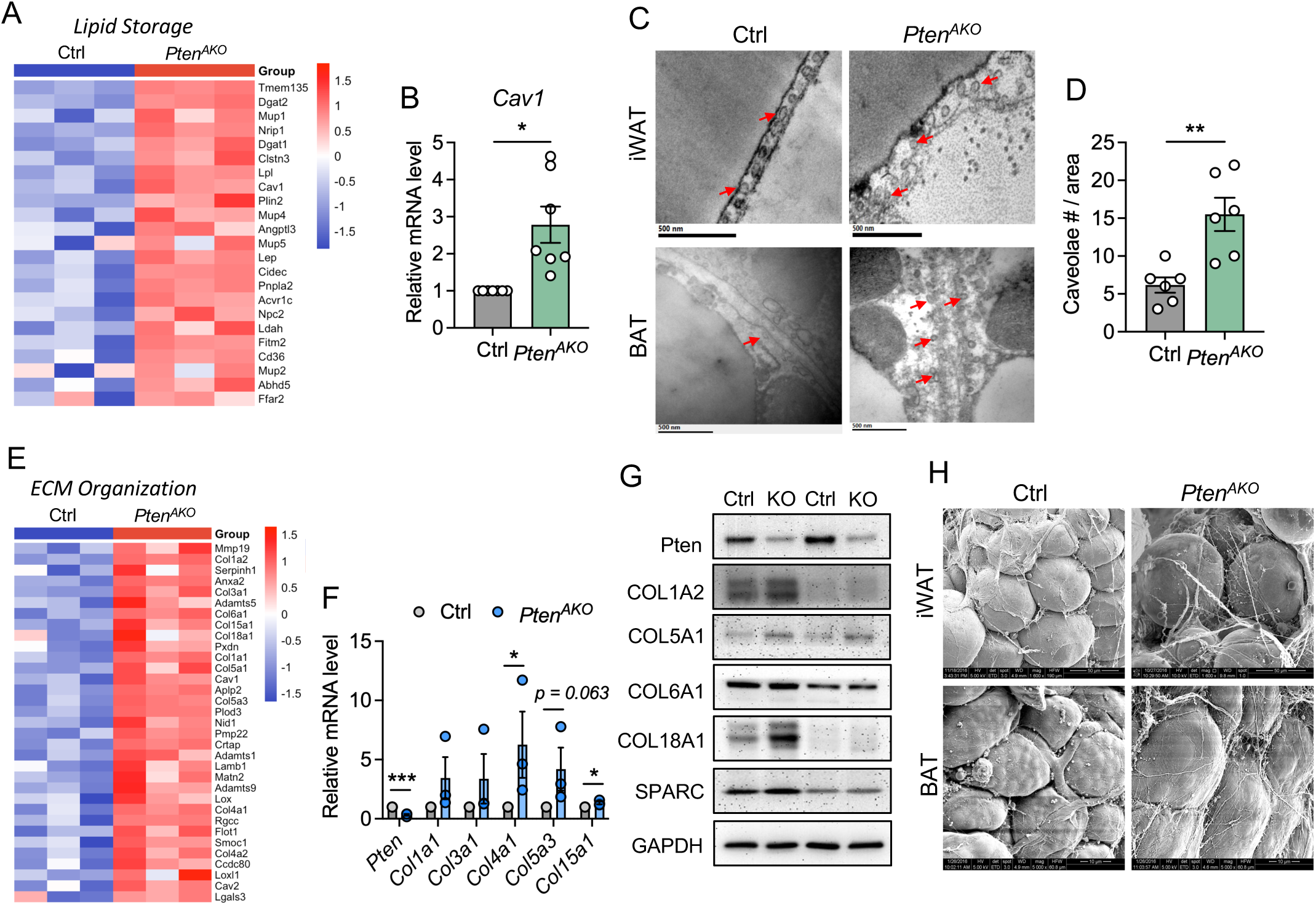
Remodeling of caveolae and ECM facilitate healthy expansion of adipose tissue. *A*: Heatmap of upregulated genes from microarray analysis related to lipid storage. *n* = 3 mice per group. *B:* qPCR analysis showing the increased expression of *Cav1* in *Pten^AKO^* iWAT. *n* = 7 mice per group. *C*: Transmission electron microscopy (TEM) imaging showing caveolae structure in iWAT and BAT of of Ctrl and *Pten^AKO^* mice. Scale bar, 500 nm. *D*: Quantification of caveolae number shown in *C*. *n* = 3 mice per group, two images were analyzed per mice. *E*: Heatmap of upregulated genes from microarray analysis related to ECM organization. *n* = 3 mice per group. *F:* qPCR analysis showing the increased expression of ECM related genes in *Pten^AKO^* iWAT. *n* = 3 mice per group. *G:* Western blot analysis of Pten and ECM proteins in iWAT of Ctrl and *Pten^AKO^* mice. *H*: Scanning electron microscopy (SEM) imaging showing ECM structure in iWAT and BAT of of Ctrl and *Pten^AKO^* mice. Scale bar, iWAT (50 µm), BAT (10 µm). Data are presented as mean ± SEM. \**P* < 0.05, \*\**P* < 0.01 and \*\*\**P* < 0.001 by two-tailed, unpaired Student *t*-test.

In parallel, transcriptomic analysis revealed increased expression of genes associated with extracellular matrix organization, including *Col1a1*, Col1a2, *Col3a1*, *Col4a1*, *Col4a2*, *Col5a1*, *Col5a3*, *Col15a1*, and *Col18a1* (Figure 7E). qPCR analysis further confirmed increased expression of *Col4a1* and *Col15a1* in *Pten^AKO^*iWAT, while *Col1a1*, *Col3a1* and *Col5a3* showed an increasing trend (Figure 7F). Western blot analysis also demonstrated increased protein levels of multiple collagen components, including COL1A2, COL5A1, COL6A1, and COL18A1, as well as SPARC, a matricellular protein important for extracellular matrix organization (Figure 7G). To assess whether these molecular changes were associated with structural remodeling of adipose tissue, we performed scanning electron microscopy (SEM) analysis of control and *Pten^AKO^* BAT and iWAT. Compared with controls, both *Pten^AKO^*BAT and iWAT exhibited more prominent extracellular matrix structures, with thicker and more abundant fibers observed between adipocytes (Figure 7H). Together, these findings indicate that adipocyte Pten deficiency might promote coordinated remodeling of the plasma membrane and extracellular matrix, which may facilitate healthy adipose tissue expansion and support enhanced lipid storage capacity.

## DISCUSSION

In this study, we demonstrate that adipocyte-specific deletion of *Pten* promotes robust adipose tissue expansion while improving systemic metabolic health. *Pten^AKO^* mice exhibited increased adiposity and marked adipocyte hypertrophy under both chow and HFD feeding conditions, yet this expansion was associated with lower circulating insulin, improved insulin sensitivity, reduced fasting glucose, and protection from hepatic lipid accumulation. These metabolic improvements occurred despite enlarged adipocytes and lipid-enriched BAT, indicating that adipocyte hypertrophy per se is not necessarily detrimental when adipose tissue retains sufficient lipid-buffering capacity and functional integrity [31, 32]. Mechanistically, Pten deficiency activated AKT signaling in adipose tissues, preserved mitochondrial protein content and structure, enhanced transcriptional programs linked to adipogenesis, lipid metabolism, insulin responsiveness, and lipid storage, and suppressed inflammatory and immune-response pathways. Together, these findings support a model in which adipocyte PTEN acts as an intrinsic brake on healthy adipose tissue expansion, and that releasing this brake allows adipose tissue to safely store excess lipid and protect other metabolic organs from lipotoxic stress.

Pten is a well-established negative regulator of the PI3K-AKT pathway, and previous studies have shown that mature-adipocyte *Pten* deletion enhances adipocyte insulin signaling and improves whole-body glucose metabolism [22–24]. Our findings are consistent with these studies but further highlight the importance of adipose depot context. In our constitutive *Adipoq*-Cre model, *Pten* deletion led to a particularly strong expansion of subcutaneous depots, including iWAT and aWAT, whereas the effect on gWAT was more modest under chow feeding and largely absent after prolonged HFD feeding. This depot-selective response may be important for the improved metabolic phenotype. Subcutaneous adipose tissue is generally considered more metabolically protective than visceral fat because it provides a safer lipid storage site and limits ectopic lipid deposition in liver, muscle, and other organs [10, 11]. Indeed, prior work has proposed that healthy expansion of subcutaneous adipose tissue can defend against caloric overload and lipotoxicity, whereas restricted subcutaneous expandability promotes lipid spillover and insulin resistance [11, 33]. Thus, the preferential expansion of subcutaneous adipose tissue in *Pten^AKO^* mice may be a key mechanism by which adipocyte Pten deficiency uncouples increased adiposity from metabolic dysfunction.

An interesting observation in this study is that BAT from *Pten^AKO^*mice displayed increased lipid accumulation and a whitening-like morphology, yet retained molecular and structural features of functional brown adipocytes. Western blot analysis showed that UCP1 and mitochondrial OXPHOS proteins were largely maintained, PGC1α was increased, and TEM analysis revealed preserved mitochondrial morphology and cristae structure. These findings suggest that lipid accumulation in BAT does not necessarily indicate loss of brown adipocyte identity or mitochondrial integrity. Because BAT mass was substantially increased in *Pten^AKO^* mice, the total BAT mitochondrial and thermogenic capacity at the depot level may be maintained, even if individual brown adipocytes store more lipid. This interpretation is consistent with the concept that BAT function depends not only on cellular lipid content but also on mitochondrial abundance, UCP1 expression, and total tissue mass [34–36]. Nevertheless, direct physiological assays, such as cold tolerance, energy expenditure, and β3-adrenergic stimulation, will be important in future studies to determine whether BAT thermogenic function is fully preserved or functionally compensated by increased BAT mass.

Obesity-associated insulin resistance is often linked to adipose inflammation, macrophage infiltration, crown-like structures, and lipotoxicity [3, 4, 37]. Our data provide evidence that these pathological features are not simply a consequence of increased fat mass or adipocyte size, but may instead reflect a failure of adipocytes to safely handle lipid overload [38]. In *Pten^AKO^* mice, adipocytes became markedly hypertrophic, yet transcriptomic analysis and qPCR validation revealed suppression of immune-response pathways, including T cell activation, chemotaxis, interferon responses, TNFα signaling, and inflammatory response pathways. Under HFD feeding, *Pten^AKO^*mice also showed reduced hepatic steatosis and improved insulin sensitivity, supporting the idea that enhanced adipocyte lipid-buffering capacity protects against systemic lipotoxicity. These findings are consistent with prior inducible adipocyte *Pten* knockout studies showing reduced adipose inflammation, reduced crown-like structures, and reduced hepatic lipid accumulation when adipocyte insulin signaling is preserved [23]. Therefore, our results support a model in which adipose inflammation is at least partly secondary to adipocyte stress, lipid spillover, and cell injury [39]. When adipocytes can expand and store lipid efficiently, immune cell recruitment and inflammatory remodeling may be minimized, thereby preserving systemic insulin sensitivity.

Our study suggests that adipocyte lipid storage capacity may be greater than previously appreciated when plasma membrane remodeling is coordinated. One mechanism controlling the upper limit of lipid storage may involve CAV1-dependent caveolae formation [40, 41]. We found increased *Cav1* expression and caveolae density along the plasma membrane of *Pten^AKO^*adipocytes. CAV1 is the structural component of caveolae, which regulate lipid trafficking, lipid droplet organization, and membrane mechanics [42–44]. Consistent with this role, Cav1-null mice exhibit reduced adipose mass, abnormal adipocyte morphology, and hypertriglyceridemia, indicating impaired adipose lipid storage [28]. Increasing CAV1 expression and caveolae density enhances the ability of adipocytes to accommodate larger lipid droplets [43]. Caveolae can flatten and disassemble under mechanical stress, serving as a membrane reservoir that buffers increases in surface tension [45]. Recent *in vivo* studies further demonstrate that caveolar reorganization supports adipocyte expansion and mechanical adaptability [44]. Human CAV1 deficiency causes congenital generalized lipodystrophy, supporting an essential role for caveolae in adipose lipid storage [46]. Thus, increased caveolae abundance may enable extreme adipocyte hypertrophy while limiting membrane stress, lipid spillover, and cellular dysfunction.

In parallel, *Pten^AKO^* iWAT showed increased expression of ECM-related genes, elevated collagen and SPARC protein levels, and more prominent matrix structures between adipocytes. Dynamic ECM remodeling is essential for adipose tissue development and expansion [47–50]. Indeed, MT1-MMP/MMP14-mediated pericellular collagen degradation is required to reduce collagen rigidity and permit adipocyte maturation within a three-dimensional matrix [51]. However, ECM accumulation can become maladaptive when matrix deposition exceeds degradation and produces a rigid, fibrotic environment. HIF-1α-driven adipose fibrosis has been linked to local inflammation and insulin resistance [52], whereas loss of collagen VI in obese mice relieved mechanical constraints, permitting adipocyte enlargement with reduced cellular stress and improved systemic metabolism [53]. The effects of ECM remodeling may also depend on the stage of adipose expansion. MMP14 activation during early obesity reduced fibrosis and facilitated metabolically healthy expansion, whereas its activation in established obesity generated collagen VI-derived endotrophin and promoted fibrosis, inflammation, and insulin resistance [54]. More recent biomechanical studies further indicate that the metabolic consequences of ECM remodeling depend on matrix composition, turnover, cross-linking, vascularization, and tissue compliance rather than ECM abundance alone [55]. In *Pten^AKO^* iWAT, increased ECM components occurred together with enhanced lipid storage, reduced inflammation, and improved insulin sensitivity. We therefore propose that these changes represent coordinated adaptive remodeling that provides structural support while preserving sufficient tissue compliance for adipocyte expansion. Nevertheless, direct measurements of ECM turnover, cross-linking, and stiffness will be needed to establish its causal contribution.

## MATERIALS AND METHODS

### Animal

All mouse strains were obtained from Jackson Laboratory (Bar Harbor, ME) under the following stock numbers: *Adipoq-Cre* (#010803) and *Pten^f/f^* (#006440). Mice were genotyped by PCR of ear DNA using genotyping protocols described by the supplier. The genotypes of experimental KO and associated control animals are as follows: *Pten^AKO^*(*Adipoq-Cre*; *Pten^f/f^*) and control (*Pten^f/f^*). Mice were housed and maintained in the animal facility with free access to water and standard rodent chow food (2018 Teklad Global 18% Protein Rodent Diets) or high-fat diet (60% Kcal from fat, TD.06414) and were housed under 12-h light/dark cycle, at 22 °C, and 45% humidity on average. For all animal-based experiments, at least three pairs of sex-matched littermates at the age of 2–6 months were used if not stated differently. For food intake study, mice were single housed in a cage for a week. Mice were anesthetized with 3% isoflurane followed by euthanasia by cervical dislocation. All procedures involving mice were approved by the Purdue University Animal Care and Use Committee.

### Body composition assessment

Body composition was measured in ad-lib fed conscious mice using an EchoMRI 3-in-1 system nuclear magnetic resonance spectrometer (EchoMedical Systems). The instrument was calibrated on each day of use with manufacturer-supplied canola-oil and phantom standards. Mice were weighed immediately before scanning. Then each mouse was placed in a ventilated, size-appropriate plastic restraint tube and scanned. Outputs included fat mass and lean mass.

### Blood glucose measurements

For glucose-tolerance test (GTT), mice were given an intraperitoneal (i.p.) injection of 100 mg/ml D-glucose (2 g/kg body weight for chow diet-fed mice and 1 g/kg body weight for HFD-fed mice) after overnight fasting, and tail blood glucose concentrations were measured by a glucometer (Accu-Check Active, Roche). For insulin-tolerance test (ITT), mice were fasted for 6 h before i.p. injection of insulin (Sigma Aldrich, cat#I6634) at the dosage of 0.75 U/kg body weight for chow diet-fed mice and 1.25 U/kg body weight for HFD-fed mice, and tail blood glucose concentrations were monitored. For both GTT and ITT, each mouse was singly caged with blinded cage numbers and random orders. Glucose excursion and insulin-stimulated glucose clearance were quantified by calculating the area under the curve (AUC) and area above the curve (AAC), respectively.

### Measurement of serological parameters

Serum levels of non-esterified fatty acids (NEFA), cholesterol, high-density lipoprotein (HDL), low-density lipoprotein (LDL), and triglycerides (TG) were examined by standard kits from Randox Laboratories.

### Transmission electron microscopy (TEM)

TEM analysis of adipose tissues was performed as described previously [56]. In brief, iBAT and iWAT were dissected, cut into 1 mm^3^ blocks, and fixed immediately in the fixative buffer (2.5% glutaraldehyde, 1.5% paraformaldehyde (PFA) in 0.1 M cacodylate buffer). Samples were rinsed in deionized water followed by fixation in 2% osmium tetroxide for 1 h. Then, the samples were washed in deionized water, followed by fixation in 1% uranyl acetate for 15 min. The samples were then dehydrated with a series of graded ethanol followed by dehydration in acetonitrile and embedded in epoxy resin (EMbed 812: DDSA: NMA 5:4:2; 0.22 DMP-30). Ultrathin sections were cut at 70 nm and stained with uranyl acetate and lead citrate. Stained sections were examined under Tecnai T12 transmission electron microscope attached with a Gatan imaging system.

### Scanning electron microscopy (SEM)

Adipose tissue morphology was examined by scanning electron microscopy (SEM). Briefly, adipose tissue samples were dissected, fixed in the fixative buffer (2.5% glutaraldehyde, 1.5% PFA in 0.1 M cacodylate buffer), processed through standard dehydration and drying procedures, and mounted on SEM stubs. Samples were then sputter-coated with a conductive metal layer prior to imaging. SEM imaging was performed using a Teneo VolumeScope field emission gun scanning electron microscope (Teneo VolumeScope FEG) at the Purdue Electron Microscopy Center. Images were acquired at appropriate accelerating voltage and magnification to visualize adipose tissue surface morphology and cellular architecture.

### H&E staining and immunohistochemistry staining

Adipose tissues were dissected and fixed in 4% PFA for 24 h at 4 °C. The tissues were dehydrated with a series of graded ethanol, cleared with xylene, and embedded into paraffin. The samples were cut into 4-μm thick slices, deparaffinized, and rehydrated using xylene, ethanol, and water by standard methods. For H&E staining, sections were stained with hematoxylin for 15 min, then rinsed in running tap water and stained with eosin for 1 min and 30 min for BAT and WAT samples, respectively. For immunohistochemistry staining, antigen retrieval was performed by submerging slides in 0.01 M sodium citrate (pH 6.0) and heated to 96 °C for 20 min in a laboratory microwave (PELCO). Slides were incubated with 3% hydrogen peroxide and 2.5% normal horse serum (S-2012, Vector), followed by incubation with rabbit polyclonal anti-UCP1 primary antibody (Abcam, cat#ab10983) diluted 1:200 in 2.5% normal horse serum (Vector, cat# S-2012) for 60 min. Signals were detected with a rabbit IgG horseradish peroxidase (HRP)-conjugated secondary antibody (Jackson ImmunoResearch, cat#111-035-003). Labeling was visualized with 3,3′-diaminobenzidine (DAB) (Acros Organics, cat#112090050) as the chromogen (Vector, cat#SK-4105). All the images were captured using a Leica DM6000B microscope. The images shown are representative results of at least three biological replicates.

### Oil Red O staining

Tissue sections were fixed with 4% PFA for 15 min at RT. After wash with PBS, the slides were stained using the Oil red O working solutions (6 mL 5 g/L Oil red O stock solution in isopropanol, 4 mL ddH_2_O, filtered for three times) for 30 min. After staining, the slides were washed with 60% isopropanol and imaged.

### Bodipy staining

Neutral lipid accumulation on tissue sections was assessed by BODIPY 493/503 staining. Briefly, tissue sections were fixed with 4% PFA for 15 min at RT. After wash with PBS, the slides were incubated with BODIPY 493/503 working solution, defined as 1 µg/mL BODIPY 493/503 (Invitrogen, cat#D3922) diluted in PBS, for 15–30 minutes at RT in the dark. After washing, nuclei were counterstained with DAPI, and samples were mounted and imaged using Leica DM6000B microscope.

### Protein extraction and immunoblotting analysis

Total protein was isolated from cells using RIPA buffer containing 25 mM Tris-HCl (pH 8.0), 150 mM NaCl, 1 mM EDTA, 0.5% NP-40, 0.5% sodium deoxycholate, and 0.1% SDS. Protein concentrations were determined using Pierce BCA Protein Assay Reagent. Proteins were separated by SDS-PAGE, transferred to a polyvinylidene fluoride membrane (Millipore Corporation), blocked in 5% fat-free milk for 1 h at RT, and then incubated with primary antibodies in 5% milk overnight at 4 °C. The membrane was then incubated with secondary antibody for 1 h at RT. Antibodies used for western blot analysis were listed in Supplementary Table 1. Immunodetection was performed using enhanced chemiluminescence western blotting substrate (Santa Cruz Biotechnology, cat#sc-2048) and detected with a FluorChem R System (ProteinSimple). The results shown in the figures are representative results from at least three independent experiments.

### Total RNA extraction and real-time PCR

Total RNA was extracted from cultured cells or mouse tissues using TRIzol reagent (Thermo Fisher Scientific, cat#15596018) according to the manufacturer’s instructions. RNA concentration and purity were assessed spectrophotometrically. Complementary DNA (cDNA) was synthesized from total RNA using random primers with M-MLV reverse transcriptase (Invitrogen, cat#28-025-021). Quantitative real-time PCR (qRT-PCR) was performed using SYBR Green Master Mix (Roche, cat#4913850001) on a LightCycler 96 Real-Time PCR System (Roche). Relative gene expression levels were calculated using the 2^⁻ΔΔCt^ method and normalized to the indicated housekeeping genes [57]. Primer sequences are listed in Supplemental Table 2.

### Microarray and GSEA analysis

Microarray was performed as described in published protocol [58, 59]. RNA was extracted from adipose tissues and gene expression was analyzed by microarray with Agilent SurePrint G3 Mouse GE 8 X 60 K chip (Agilent, cat#G4858A). The list of significantly changed genes with a fold change ≥1.5-fold was used for Gene Ontology (GO) pathway analysis. Gene set enrichment analysis (GSEA) was conducted as described previously [58, 59].

### Statistical analysis

All quantitative data are presented as mean ± standard error of the mean (SEM) from at least three independent biological replicates except two replicates for microarray analysis. Statistical significance between two groups was determined using a two-tailed Student’s *t*-test. A *P* value < 0.05 was considered statistically significant.

## Supporting information

Supplementary Datasheet 1

Supplementary Datasheet 2

Supplementary Datasheet 3

## DATA AVAILABILITY STATEMENT

All data reported in this manuscript will be shared by the lead contact upon request. No original code was reported that needed to reanalyze the data generated by this study. The microarray data generated in this study have been provided as supplementary datasheet. Any additional information required to reanalyze the data reported in this manuscript is available from the corresponding author upon request.

## AUTHOR CONTRIBUTIONS

**Yumei Zhou**: Data curation; Formal analysis; Investigation; Visualization; Methodology; Writing—original draft. **Yubo Wang**: Data curation; Formal analysis. **Jessica E. Meerson**: Data curation; Formal analysis. **Zhiyong Cheng**: Writing—review and editing. **Shihuan Kuang**: Resources; Supervision; Funding acquisition; Writing—review and editing. **Feng Yue**: Conceptualization; Data curation; Software; Formal analysis; Supervision; Funding acquisition; Project administration; Validation; Investigation; Visualization; Methodology; Writing—review and editing.

## ACKNOWLEDGMENTS

This work was supported by grants from the National Institutes of Health NIH-R01DK136722 to F.Y., NIH-R01DK132819, NIH-R01AR078695, and NIH-R01AR079235 to S.K., and the University of Florida Research Startups to F.Y. We thank Dr. Jennifer Freeman and Sara Wirbisky at Purdue University for helping with microarray analysis, Dr. Naagarajan Narayanan and Dr. Meng Deng for assisting the TEM and SEM analysis, and the other Yue and Kuang lab members for their discussion and assistance. We are grateful to Purdue Electron Microscopy Facility for assistance on TEM and SEM analysis.

## REFERENCES

1. Sakers A, De Siqueira MK, Seale P, Villanueva CJ, "Adipose-tissue plasticity in health and disease," Cell 185 (2022): 419–446.

2. Asterholm IW, Tao C, Morley TS, Wang QA, Delgado-Lopez F, Wang ZV, et al., "Adipocyte inflammation is essential for healthy adipose tissue expansion and remodeling," Cell metabolism 20 (2014): 103–118.

3. Santoro A, McGraw TE, Kahn BB, "Insulin action in adipocytes, adipose remodeling, and systemic effects," Cell metabolism 33 (2021): 748–757.

4. Wu H, Ballantyne CM, "Metabolic inflammation and insulin resistance in obesity," Circulation research 126 (2020): 1549–1564.

5. Shoelson SE, Lee J, Goldfine AB, "Inflammation and insulin resistance," The Journal of clinical investigation 116 (2006): 1793–1801.

6. Longo M, Zatterale F, Naderi J, Parrillo L, Formisano P, Raciti GA, et al., "Adipose tissue dysfunction as determinant of obesity-associated metabolic complications," International journal of molecular sciences 20 (2019): 2358.

7. Morigny P, Boucher J, Arner P, Langin D, "Lipid and glucose metabolism in white adipocytes: pathways, dysfunction and therapeutics," Nature Reviews Endocrinology 17 (2021): 276–295.

8. Hagberg CE, Spalding KL, "White adipocyte dysfunction and obesity-associated pathologies in humans," Nature Reviews Molecular Cell Biology 25 (2024): 270–289.

9. Arner P, Bernard S, Salehpour M, Possnert G, Liebl J, Steier P, et al., "Dynamics of human adipose lipid turnover in health and metabolic disease," Nature 478 (2011): 110–113.

10. Blüher M, "Metabolically healthy obesity," Endocrine reviews 41 (2020): bnaa004.

11. Schulze MB, Stefan N, "Metabolically healthy obesity: from epidemiology and mechanisms to clinical implications," Nature Reviews Endocrinology 20 (2024): 633–646.

12. Iacobini C, Pugliese G, Fantauzzi CB, Federici M, Menini S, "Metabolically healthy versus metabolically unhealthy obesity," Metabolism 92 (2019): 51–60.

13. Smith GI, Mittendorfer B, Klein S, "Metabolically healthy obesity: facts and fantasies," The Journal of clinical investigation 129 (2019): 3978–3989.

14. Vishvanath L, Gupta RK, "Contribution of adipogenesis to healthy adipose tissue expansion in obesity," The Journal of clinical investigation 129 (2019): 4022–4031.

15. Kolb H, "Obese visceral fat tissue inflammation: from protective to detrimental?," BMC medicine 20 (2022): 494.

16. Veilleux A, Caron-Jobin M, Noël S, Laberge PY, Tchernof A, "Visceral adipocyte hypertrophy is associated with dyslipidemia independent of body composition and fat distribution in women," Diabetes 60 (2011): 1504–1511.

17. Verboven K, Wouters K, Gaens K, Hansen D, Bijnen M, Wetzels S, et al., "Abdominal subcutaneous and visceral adipocyte size, lipolysis and inflammation relate to insulin resistance in male obese humans," Scientific reports 8 (2018): 4677.

18. Chen C-Y, Chen J, He L, Stiles BL, "PTEN: tumor suppressor and metabolic regulator," Frontiers in endocrinology 9 (2018): 338.

19. Song MS, Salmena L, Pandolfi PP, "The functions and regulation of the PTEN tumour suppressor," Nature reviews Molecular cell biology 13 (2012): 283–296.

20. Pal A, Barber TM, Van de Bunt M, Rudge SA, Zhang Q, Lachlan KL, et al., "PTEN mutations as a cause of constitutive insulin sensitivity and obesity," New England Journal of Medicine 367 (2012): 1002–1011.

21. Ortega-Molina A, Efeyan A, Lopez-Guadamillas E, Muñoz-Martin M, Gómez-López G, Cañamero M, et al., "Pten positively regulates brown adipose function, energy expenditure, and longevity," Cell metabolism 15 (2012): 382–394.

22. Kurlawalla-Martinez C, Stiles B, Wang Y, Devaskar SU, Kahn BB, Wu H, "Insulin hypersensitivity and resistance to streptozotocin-induced diabetes in mice lacking PTEN in adipose tissue," Mol Cell Biol 25 (2005): 2498–2510.

23. Morley TS, Xia JY, Scherer PE, "Selective enhancement of insulin sensitivity in the mature adipocyte is sufficient for systemic metabolic improvements," Nature communications 6 (2015): 7906.

24. Huang W, Queen NJ, McMurphy TB, Ali S, Cao L, "Adipose PTEN regulates adult adipose tissue homeostasis and redistribution via a PTEN-leptin-sympathetic loop," Molecular metabolism 30 (2019): 48–60.

25. Kirstein AS, Kehr S, Nebe M, Hanschkow M, Barth LA, Lorenz J, et al., "PTEN regulates adipose progenitor cell growth, differentiation, and replicative aging," Journal of Biological Chemistry 297 (2021): 100968.

26. Lee KY, Russell SJ, Ussar S, Boucher J, Vernochet C, Mori MA, et al., "Lessons on conditional gene targeting in mouse adipose tissue," Diabetes 62 (2013): 864–874.

27. Zhang C, Ye M, Melikov K, Yang D, Vale GDd, McDonald J, et al., "CLSTN3B promotes lipid droplet maturation and lipid storage in mouse adipocytes," Nature communications 15 (2024): 9475.

28. Razani B, Combs TP, Wang XB, Frank PG, Park DS, Russell RG, et al., "Caveolin-1-deficient mice are lean, resistant to diet-induced obesity, and show hypertriglyceridemia with adipocyte abnormalities," Journal of Biological Chemistry 277 (2002): 8635–8647.

29. Pilch PF, Liu L, "Fat caves: caveolae, lipid trafficking and lipid metabolism in adipocytes," Trends in Endocrinology & Metabolism 22 (2011): 318–324.

30. Sun K, Kusminski CM, Scherer PE, "Adipose tissue remodeling and obesity," The Journal of clinical investigation 121 (2011): 2094–2101.

31. Gustafson B, Hedjazifar S, Gogg S, Hammarstedt A, Smith U, "Insulin resistance and impaired adipogenesis," Trends in Endocrinology & Metabolism 26 (2015): 193–200.

32. Laforest S, Labrecque J, Michaud A, Cianflone K, Tchernof A, "Adipocyte size as a determinant of metabolic disease and adipose tissue dysfunction," Critical reviews in clinical laboratory sciences 52 (2015): 301–313.

33. Virtue S, Vidal-Puig A, "Adipose tissue expandability, lipotoxicity and the metabolic syndrome—an allostatic perspective," Biochimica et Biophysica Acta (BBA)-Molecular and Cell Biology of Lipids 1801 (2010): 338–349.

34. Leitner BP, Huang S, Brychta RJ, Duckworth CJ, Baskin AS, McGehee S, et al., "Mapping of human brown adipose tissue in lean and obese young men," Proceedings of the national academy of sciences 114 (2017): 8649–8654.

35. Cannon B, Nedergaard J, "Brown adipose tissue: function and physiological significance," Physiological reviews 84 (2004): 277–359.

36. Cypess AM, Cannon B, Nedergaard J, Kazak L, Chang DC, Krakoff J, et al., "Emerging debates and resolutions in brown adipose tissue research," Cell metabolism 37 (2025): 12–33.

37. Zatterale F, Longo M, Naderi J, Raciti GA, Desiderio A, Miele C, et al., "Chronic adipose tissue inflammation linking obesity to insulin resistance and type 2 diabetes," Frontiers in physiology 10 (2020): 1607.

38. Kim JI, Huh JY, Sohn JH, Choe SS, Lee YS, Lim CY, et al., "Lipid-overloaded enlarged adipocytes provoke insulin resistance independent of inflammation," Molecular and cellular biology 35 (2015): 1686–1699.

39. Saltiel AR, Olefsky JM, "Inflammatory mechanisms linking obesity and metabolic disease," The Journal of clinical investigation 127 (2017): 1–4.

40. Tol MJ, Shimanaka Y, Bedard AH, Sapia J, Cui L, Colaço-Gaspar M, et al., "Dietary control of peripheral adipose storage capacity through membrane lipid remodelling," Nature Metabolism 7 (2025): 1424–1442.

41. Haczeyni F, Bell-Anderson KS, Farrell G, "Causes and mechanisms of adipocyte enlargement and adipose expansion," Obesity reviews 19 (2018): 406–420.

42. Cohen AW, Hnasko R, Schubert W, Lisanti MP, "Role of caveolae and caveolins in health and disease," Physiological reviews 84 (2004): 1341–1379.

43. Briand N, Prado C, Mabilleau G, Lasnier F, Le Lièpvre X, Covington JD, et al., "Caveolin-1 expression and cavin stability regulate caveolae dynamics in adipocyte lipid store fluctuation," Diabetes 63 (2014): 4032–4044.

44. Aboy-Pardal MC, Guadamillas MC, Guerrero CR, Català-Montoro M, Toledano-Donado M, Terrés-Domínguez S, et al., "Plasma membrane remodeling determines adipocyte expansion and mechanical adaptability," Nature Communications 15 (2024): 10102.

45. Sinha B, Köster D, Ruez R, Gonnord P, Bastiani M, Abankwa D, et al., "Cells respond to mechanical stress by rapid disassembly of caveolae," Cell 144 (2011): 402-413.

46. Kim CA, Delépine M, Boutet E, El Mourabit H, Le Lay S, Meier M, et al., "Association of a homozygous nonsense caveolin-1 mutation with Berardinelli-Seip congenital lipodystrophy," The Journal of Clinical Endocrinology & Metabolism 93 (2008): 1129–1134.

47. Mariman EC, Wang P, "Adipocyte extracellular matrix composition, dynamics and role in obesity," Cellular and molecular life sciences 67 (2010): 1277–1292.

48. Sun K, Tordjman J, Clément K, Scherer PE, "Fibrosis and adipose tissue dysfunction," Cell metabolism 18 (2013): 470–477.

49. Sun K, Li X, Scherer PE, "Extracellular matrix (ECM) and fibrosis in adipose tissue: overview and perspectives," Comprehensive Physiology 13 (2023): 4387–4407.

50. Ruiz-Ojeda FJ, Méndez-Gutiérrez A, Aguilera CM, Plaza-Díaz J, "Extracellular matrix remodeling of adipose tissue in obesity and metabolic diseases," International journal of molecular sciences 20 (2019): 4888.

51. Chun T-H, Hotary KB, Sabeh F, Saltiel AR, Allen ED, Weiss SJ, "A pericellular collagenase directs the 3-dimensional development of white adipose tissue," Cell 125 (2006): 577–591.

52. Halberg N, Khan T, Trujillo ME, Wernstedt-Asterholm I, Attie AD, Sherwani S, et al., "Hypoxia-inducible factor 1α induces fibrosis and insulin resistance in white adipose tissue," Molecular and cellular biology 29 (2009): 4467–4483.

53. Khan T, Muise ES, Iyengar P, Wang ZV, Chandalia M, Abate N, et al., "Metabolic dysregulation and adipose tissue fibrosis: role of collagen VI," Molecular and cellular biology 29 (2009): 1575–1591.

54. Li X, Zhao Y, Chen C, Yang L, Lee H-h, Wang Z, et al., "Critical role of matrix metalloproteinase 14 in adipose tissue remodeling during obesity," Molecular and cellular biology 40 (2020): e00564-00519.

55. Unamuno X, Gomez-Ambrosi J, Becerril S, Álvarez-Cienfuegos FJ, Ramirez B, Rodriguez A, et al., "Changes in mechanical properties of adipose tissue after bariatric surgery driven by extracellular matrix remodelling and neovascularization are associated with metabolic improvements," Acta Biomaterialia 141 (2022): 264-279.

56. Qiu J, Yue F, Zhu P, Chen J, Xu F, Zhang L, et al., "FAM210A is essential for cold-induced mitochondrial remodeling in brown adipocytes," Nature Communications 14 (2023): 6344.

57. Livak KJ, Schmittgen TD, "Analysis of relative gene expression data using real-time quantitative PCR and the 2− ΔΔCT method," methods 25 (2001): 402–408.

58. Oprescu SN, Horzmann KA, Yue F, Freeman JL, Kuang S, "Microarray, IPA and GSEA analysis in mice models," Bio-protocol 8 (2018): e2999–e2999.

59. Bi P, Yue F, Sato Y, Wirbisky S, Liu W, Shan T, et al., "Stage-specific effects of Notch activation during skeletal myogenesis," Elife 5 (2016): e17355.

