## Supplementary Table 1 for "Adipocyte Pten Inhibition Improves Metabolic Health Associated with Expanded Lipid Storage Capacity and Reduced Inflammation"

### List of antibodies used in this study

| Antibody | Manufacture | Catalog # | Application |
|---|---|---|---|
| Rabbit monoclonal anti-Pten | Cell Signaling Technology | #9188S | WB (1:1000) |
| Rabbit monoclonal anti-Phospho-Akt (Ser473) | Cell Signaling Technology | #4060S | WB (1:2000) |
| Rabbit monoclonal anti-Akt (pan) | Cell Signaling Technology | #4691S | WB (1:2000) |
| Mouse monoclonal anti-PGC1a | Santa Cruz Biotechnology | sc-13067 | WB (1:1000) |
| Mouse monoclonal anti-PPARγ | Santa Cruz Biotechnology | sc-390740 | WB (1:1000) |
| Rabbit polyclonal anti-UCP1 | AbCam | ab10983 | IHC (1:200), WB (1:1000) |
| Rabbit polyclonal anti-FABP4 | Santa Cruz Biotechnology | sc-271529 | WB (1:1000) |
| Mouse monoclonal anti-COL1A2 | Santa Cruz Biotechnology | sc-393573 | WB (1:500) |
| Mouse monoclonal anti-COL5A1 | Santa Cruz Biotechnology | sc-166155 | WB (1:500) |
| Mouse monoclonal anti-COL6A1 | Santa Cruz Biotechnology | sc-377143 | WB (1:500) |
| Mouse monoclonal anti-COL18A1 | Santa Cruz Biotechnology | sc-32720 | WB (1:500) |
| Mouse monoclonal anti-SPARC | Santa Cruz Biotechnology | sc-73472 | WB (1:500) |
| Mouse OxPhos Rodent WB Antibody Cocktail | Invitrogen | 45-8099 | WB (1:2000) |
| HRP AffiniPure goat anti-mouse IgG | Jackson ImmunoResearch | 115-035-003 | WB (1:10,000) |
| HRP AffiniPure goat anti-rabbit IgG | Jackson ImmunoResearch | 111-035-003 | WB (1:10,000) |
