## Supplementary Table 2 for "Adipocyte Pten Inhibition Improves Metabolic Health Associated with Expanded Lipid Storage Capacity and Reduced Inflammation"

**qPCR primer sequences**

| **Gene name** | **Sequence (5' to 3')** |
|---|---|
| *Glut4*: forward | GTGACTGGAACACTGGTCCTA |
| *Glut4*: reverse | CCAGCCACGTTGCATTGTAG |
| *Pfkfb4*: forward | AGGATGTCTTGTCGCCTCTG |
| *Pfkfb4*: reverse | CGGTACTTGTCCTGATCCCG |
| *Gyk*: forward | TGAAGAAAGCGAAATCCGTTACT |
| *Gyk*: reverse | CCCAAAGGCAGACTACAGAAG |
| *Mgat2*: forward | CGGAAGGTGCTAATCCTGACG |
| *Mgat2*: reverse | CCGATTCGTTTGAGACCCTG |
| *Lpgat1*: forward | CTGGGCTGGATTGTAGCGAAA |
| *Lpgat1*: reverse | GTCCTTCGATATACCAGAAGCG |
| *Agpat2*: forward | CAGCCAGGTTCTACGCCAAG |
| *Agpat2*: reverse | TGATGCTCATGTTATCCACGGT |
| *Slc27a2*: forward | TCCTCCAAGATGTGCGGTACT |
| *Slc27a2*: reverse | TAGGTGAGCGTCTCGTCTCG |
| *Plin2*: forward | GACCTTGTGTCCTCCGCTTAT |
| *Plin2*: reverse | CAACCGCAATTTGTGGCTC |
| *Cd68*: forward | TGTCTGATCTTGCTAGGACCG |
| *Cd68*: reverse | GAGAGTAACGGCCTTTTTGTGA |
| *Tnfa*: forward | CTGAACTTCGGGGTGATCGG |
| *Tnfa*: reverse | GGCTTGTCACTCGAATTTTGAGA |
| *Il1b*: forward | GACCTTGTGTCCTCCGCTTAT |
| *Il1b*: reverse | AGCAGCCCTTCATCTTTTGG |
| *Cav1*: forward | ATGTCTGGGGGCAAATACGTG |
| *Cav1*: reverse | CGCGTCATACACTTGCTTCT |
| *Il1b*: forward | GACCTTGTGTCCTCCGCTTAT |
| *Il1b*: reverse | CAACCGCAATTTGTGGCTC |
| *Col1a1*: forward | GCTCCTCTTAGGGGCCACT |
| *Col1a1*: reverse | CCACGTCTCACCATTGGGG |
| *Col3a1*: forward | CTGTAACATGGAAACTGGGGAAA |
| *Col3a1*: reverse | CCATAGCTGAACTGAAAACCACC |
| *Col4a1*: forward | CTGGCACAAAAGGGACGAG |
| *Col4a1*: reverse | ACGTGGCCGAGAATTTCACC |
| *Col5a3*: forward | CGGGGTACTCCTGGTCCTAC |
| *Col5a3*: reverse | GCATCCCTACTTCCCCCTTG |
| *Col15a1*: forward | TTGACGGTCGGGATGTGATG |
| *Col15a1*: reverse | TACTGCCATGTCCGTGGTTC |
| *Gapdh*: forward | AGGTCGGTGTGAACGGATTTG |
| *Gapdh*: reverse | TGTAGACCATGTAGTTGAGGTCA |
